# Diversified RACE Sampling on Data Streams Applied to Metagenomic Sequence Analysis

**DOI:** 10.1101/852889

**Authors:** Benjamin Coleman, Benito Geordie, Li Chou, R. A. Leo Elworth, Todd J. Treangen, Anshumali Shrivastava

## Abstract

The rise of whole-genome shotgun sequencing (WGS) has enabled numerous breakthroughs in large-scale comparative genomics research. However, the size of genomic datasets has grown exponentially over the last few years, leading to new challenges for traditional streaming algorithms. Modern petabyte-sized genomic datasets are difficult to process because they are delivered by high-throughput data streams and are difficult to store. As a result, many traditional streaming problems are becoming increasingly relevant. One such problem is the task of constructing a maximally diverse sample over a data stream. In this regime, complex sampling procedures are not possible due to the overwhelming data generation rate. In theory, the best diversity sampling methods are based on a simple greedy algorithm that compares the current sequence with a large pool of sampled sequences and decides whether to accept or reject the sequence. While these methods are elegant and optimal, they are largely confined to the theoretical realm because the greedy procedure is too slow in practice. While there are many methods to identify common elements in data streams efficiently, fast and memory-efficient diversity sampling remains a challenging and fundamental data streaming problem with few satisfactory solutions. In this work, we bridge the gap with RACE sampling, an online algorithm for diversified sampling. Unlike random sampling, which samples uniformly, RACE selectively accepts samples from streams that lead to higher sequence diversity. At the same time, RACE is as computationally efficient as random sampling and avoids pairwise similarity comparisons between sequences. At the heart of RACE lies an efficient lookup array constructed using locality-sensitive hashing (LSH). Our theory indicates that an accept/reject procedure based on LSH lookups is sufficient to obtain a highly diverse subsample. We provide rigorous theoretical guarantees for well-known biodiversity indices and show that RACE can nearly double the Shannon and Simpson indices of a genetic sample in practice, all while using the same resources as random sampling. We also compare RACE against Diginorm and coreset-based diversity sampling methods and find that RACE is faster and more memory efficient. Our algorithm is straightforward to implement, easy to parallelize, and fast enough to keep pace with the overwhelming data generation rates. We expect that as DNA sequence data streams become more mainstream and faster, RACE will become an essential component for many applications.^1^

## 1 Introduction

DNA sequencing plays an essential role in advancing biological research and in various applications such as virology and medical diagnosis. In particular, fast and efficient analysis of DNA sequences improves the ability of researchers to detect and catalog diverse sets of organisms and species [25]. Large scale genomic repositories such as the European Nucleotide Archive (ENA) [15] and the NCBI Short Read Archive (SRA)[16] are the product of ambitious experimental efforts to sequence as many organisms as possible. These datasets are important for a wide array of applications to human health, including pathogen surveillance [26], antibiotic resistance detection [14], and cancer genomics [21]. However, the size of these archives now rivals the largest datasets at web-scale companies and government agencies. The SRA currently contains over 30 million FASTQ files with over 26 petabytes, and the ERA contains one-fifth of a petabyte of bacterial and viral DNA alone. Historically, the total size of sequencing data has doubled every 2 years. Furthermore, the rate of exponential growth is likely to increase thanks to recent developments in sequencing hardware.

Currently, it is possible to generate very large quantities of sequencing data at extremely fast data rates. For instance, the PromethION system from Oxford Nanopore Technologies has a data throughput rate of over 4 terabytes per day. Plans are underway to parallelize the data collection process and increase the data rate even further. Although very recent base calling algorithms, such as Guppy, can potentially keep up with the data stream, researchers are generating datasets much faster than they can be transmitted to online archives. These datasets are large enough that it is a practical challenge to store them locally. As a result, large-scale genome processing, transmission and storage pose problems from an algorithmic and systems perspective. Based on historical trends and recent hardware, it is likely that storage and transmission of the complete data will soon become infeasible. To cope with the onslaught of data, the algorithms community will need to develop streaming algorithms that are appropriate for sequencing data.

In spite of recent methods designed to handle the data deluge [25], new methods are needed that focus on data compression, sampling, and summarization to facilitate faster algorithms and practical systems at such scale. Since sequence datasets contain highly redundant information, we may be able to represent the same scientifically relevant information in much less space by selectively sampling sequences from the data stream. It is clear that the sampling process cannot require more than one pass through these overwhelmingly large datasets. The most popular approach for sampling on a data stream is reservoir sampling, which returns a uniform random sample from the stream after one pass through the data. However, random samples lose rare information. In fields such as metagenomic studies, where we wish to identify all organisms present in a population, we often need to preserve rare sequence. Most sequences will belong to common organisms with large sub-populations, but relatively few sequences will correspond to rare organisms. Therefore, the probability of retaining information from these rare organisms under random sampling is suboptimal. While this issue can be resolved by storing a very large number of samples, this is undesirable because it increases the amount of genetic information passed to computationally-expensive downstream pipelines.

Inspired by these developments, we consider the streaming diversity sampling problem. In this problem, we are given a data stream 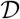 of elements (or sequences) *x*_1_, *x*_2_,…, *x_N_*, which we see one element at a time. The task is to construct a diverse sample 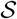 of 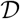 using limited computational resources. In particular, we expect the following three properties to be satisfied by the sampling algorithm:

1. **One Pass:** It is clearly prohibitive to make a second pass over the stream. Thus, once we see *x_i_* we must immediately decide whether to accept or reject it and move on. It should be noted that this decision is oblivious to all future samples *x*_*i*+1_, *x*_*i*+2_, …, *x_N_*
2. **Efficient Computation:** An intuitive and straightforward solution is to store a buffer of elements and compare the current element *x_i_* to all the sampled elements. While this will preserve diversity, it is very slow for large sample sizes. Large numbers of comparisons are infeasible in the online setting because the data generation rate is usually much faster than a linear scan with pairwise comparisons. For datasets at the petabyte scale, this procedure is prohibitively slow.
3. **Tiny Main Memory:** There are several efficient near-neighbor based methods which can return the closest element efficiently, but they all require expensive data structures that are super-linear in the size of the data. Since these structures must reside in main memory, such methods are prohibitive. We can expect a reasonable sub-sample of peta-scale datasets to be sufficiently large that it does not fit in any existing RAM. Therefore, we cannot perform near-neighbor lookups on genetic data streams in practice.

Our aim is to perform diverse sampling while achieving the above three properties. Diversity maximization problems have been explored by the computational geometry community in the context of diverse coresets [11] and diverse near-neighbor search [2]. Diversity is usually defined geometrically. For instance, a common definition is that diversity is equal to the minimum pairwise distance between the points of the sample set. However, other diversity measures are more useful and well-known for genomics problems. In this paper we will focus on biodiversity measures such as the Simpson index and Shannon index which are popular in the bioinformatics community.

### 1.1 Related Work

One can always construct a random sample of the dataset using the popular reservoir method originally proposed by [27]. Random sampling has all three aforementioned properties but is undesirable because it discards important information. In this section, we review related ideas and their shortcomings.

#### Composable Coresets

To the best of our knowledge, the only direct solution to the diversity sampling problem is the approach proposed by [11]. This algorithm is based on the idea of *composable coresets.* Suppose we want to efficiently solve an optimization problem on a large dataset 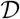. A coreset is a subset 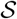 of 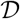 with the following property. If we solve the optimization problem on 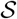, we obtain an approximate solution to the problem on 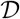. Since 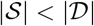, coresets enable fast processing of massive datasets with limited memory. Now suppose we construct a coreset 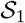 for dataset 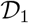 and 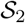 for dataset 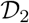. The coresets are composable if we can *merge* 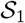 and 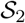 to get a coreset for the combined dataset 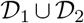. Composable coresets can solve streaming problems by breaking the stream into buffers of *M* elements. We begin by constructing a composable coreset 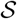 for the first *M* elements of a data stream. To process the next *M* elements, we construct another coreset and merge it with 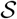 [11]. Since the cost of finding the coreset scales with *M*, the buffer size *M* is chosen to fit within computational constraints. The output is a diverse coreset of the entire data stream. Although the merging and construction operations can be costly, this approach seems to work well in practice on web-scale streaming data [1].

#### Buffer Algorithms

Another category of algorithms use a buffer to identify the most frequent elements. Two well-known algorithms are *lossy counting* (deterministic) and *sticky sampling* (probabilistic) [18]. Lossy counting records the frequency of each unique item in an array. After processing each chunk of the data stream, we decrement the counters and remove any elements with zero count values. Lossy counting reports a slight underestimate of the element frequencies but provably identifies the most common elements in the stream.

Sticky sampling is a related method for the same problem. Rather than keep a fixed-size array, sticky sampling dynamically allocates a new counter when it samples a new element. Unlike lossy counting, sticky sampling doubles the chunk size each iteration while randomly sampling over the chunk. Elements are collected from the stream with probability 1/*r*, where *r* is the chunk size. After each chunk, the count values in the buffer are decremented according to a random process. As before, elements are removed from the buffer when they have zero count value.

#### Sketching Algorithms

Sketching algorithms construct a compact (sub-linear space) data structure to summarize information from large quantities of data. The Count-Min sketch (CMS) is a data summary that can efficiently approximate how frequently an element occurs in a data stream [7]. The CMS consists of an array of counters that is randomly indexed using a hash function. When a new element *x* arrives from the stream, it is hashed to a small set of counters, which are incremented. Each counter keeps track of how many times *x* has been seen, plus the number of times another element has mapped to the same counter or had a hash collision. By randomly merging counters, the CMS provides an accurate frequency estimate for the most frequent items of a data stream. Diginorm [5], Bignorm [28] and NeatFreq [20] are sketching algorithms for genetic data normalization that use the CMS. These methods reduce the size of genetic datasets by eliminating redundant reads. At a high level, Diginorm and Bignorm use the CMS to downsample high-frequency k-mers while attempting to retain rare information.

### 1.2 Limitations: Why are existing methods insufficient?

Existing algorithms have two key limitations that prevent their application to large-scale data streams with fast data rates such as genome sequence data streams. The streaming algorithm literature is largely focused on locating frequent elements and eliminating exact duplicates in the data stream. Such methods lack robustness to even a slight perturbation in the elements. Thus, a CMS or lossy counting buffer can answer whether we have seen a particular sequence but cannot tell whether we have seen any similar sequences. Duplicate detection will avoid redundancy but may not actually increase the diversity of our sample, especially since many sequences that are otherwise identical will have small perturbations. Furthermore, the sketch size grows with the number of sequences in the stream, making duplicate detection expensive and prohibitive for long-running data streams. This is reflected in our experiments, where Diginorm [5] required very large amounts of memory.

Composable coresets are the only known streaming algorithm that can reject similar sequences, but they scale poorly with sample size. As a result, composable coresets are infeasible for practical data streams. In our experiments, we could not scale coresets to work on genetic datasets with more than 1 million reads. In this paper, we provide a significant departure from the existing line of work and propose a novel sampling algorithm. We use the statistical view of locality-sensitive hashing to bypass distance computations while still preserving diversity and robustness to perturbation.

## 2 RACE Sampling Algorithm

We propose a fast diversity sampling routine that handles high-throughput streams of sequence data. Our method constructs a diverse sample set by rejecting sequences that are similar to the collected samples. The most critical part is that we do not perform any kind of pairwise distance computation. We are able to avoid distance computations by using locality-sensitive hashing and the recently proposed RACE algorithm. Before describing our algorithm, we introduce two fundamental components of our proposal.

**Locality-Sensitive Hashing** A locality-sensitive hash (LSH) family [12] is a family of functions with the following property: Under the hash mapping, similar points have a high probability of having the same hash value.

### Definition 1.

(*R, cR, α, β*)-*sensitive hash family*

*A family* 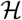 *is called* (*R, cR, α, β*)-*sensitive with respect to a distance function d*(*x, y*) *if the following properties hold for* 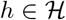 *and any two points x and y*:

– *If d*(*x, y*) ≤ *R then* 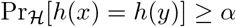
– *If d*(*x, y*) ≥ *cR then* 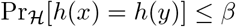

The two points *x* and *y* are said to *collide* if *h*(*x*) = *h*(*y*). We will use the notation *ρ*(*d*(*x, y*)) to refer to the collision probability 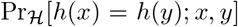. Note that *ρ* is only a function of the distance *d*(*x, y*) between *x* and *y*. Also, observe that we can increase the sensitivity of a LSH function. For any positive integer *n*, if there exists a (*R, cR, α, β*)-sensitive LSH function *h*(·) with *ρ*, then the same hash function can be independently concatenated *n* times to obtain a new hash function that is (*R, cR, α^n^, β^n^*)-sensitive with collision probability *ρ^n^*. Finally, we introduce the rehashing trick. If a LSH function *h*(·) has collision probability *ρ* and we hash the LSH values to a finite range [1, *R*] using a universal hash function, then the new hash function is locality sensitive with collision probability 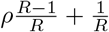.

**RACE Sketch** The RACE sketch is an array *A* of counters indexed by a LSH function. When a new data element *x* arrives from the stream, we hash *x* using *R* different LSH functions to get *R* indices. We increment the RACE counters at these indices and process the next element. The RACE sketch can also be queried to quickly determine whether we have already seen a data element similar to *x*. To do this, we take the mean of the count values at the *R* indices corresponding to *x*. The fundamental theoretical result from [17] and [6] is that this process approximates the sum of LSH collision probabilities, which is a useful geometric object known as a kernel density estimate.

### Theorem 1.

*ACE Estimator*

*Given a dataset* 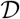 *and a LSH family* 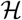, *construct an ACE array A a LSH function h*(*q*) *from* 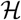. *Then for any query q*,

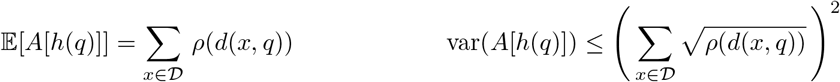

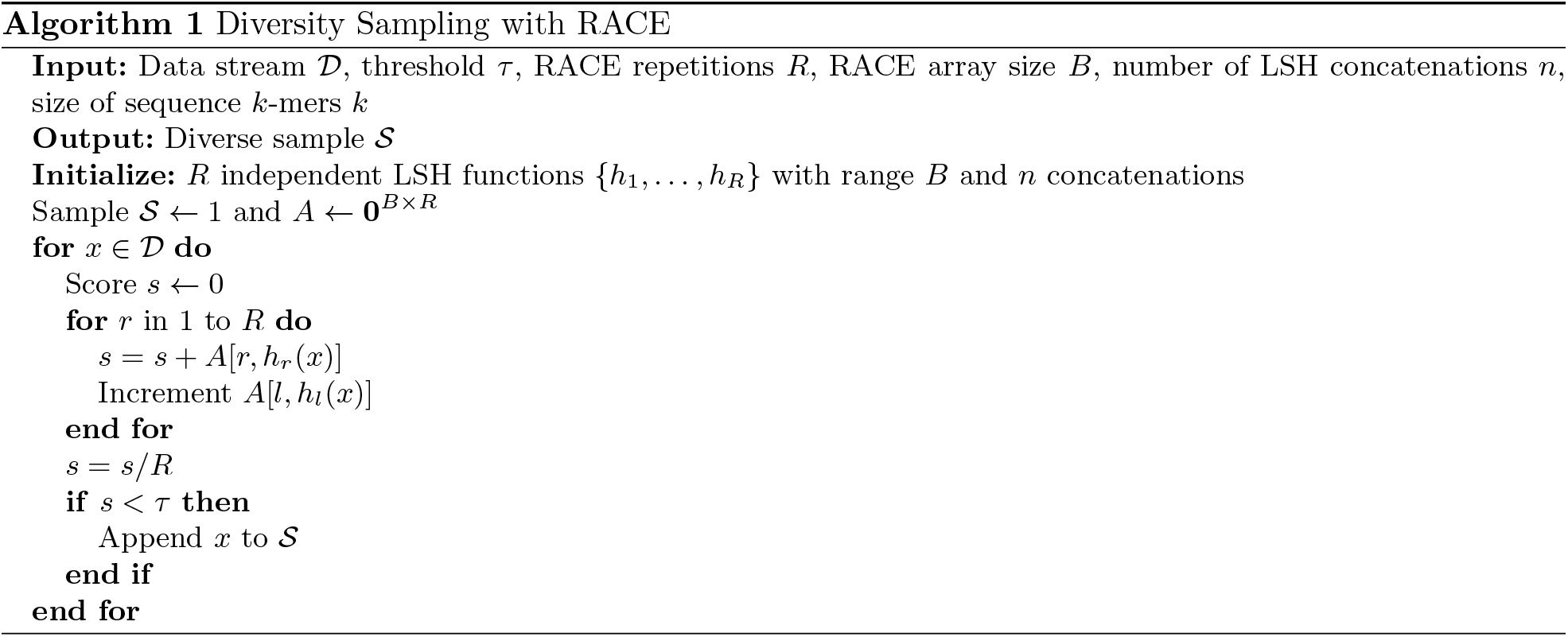

By querying several ACE array repetitions (RACE), we obtain a good estimate of 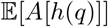. The key intuition behind RACE is that if 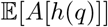 is small, then *q* is far away from most elements in the dataset since there are many low collision probabilities. Therefore, *x* is an outlier and we have not seen many similar elements.

### 2.1 Our Proposal

We propose a sampling algorithm that uses RACE to select a diverse set of sequences. When a new sequence *x* arrives from the stream, we estimate 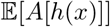, which we will refer to as the RACE score *s*(*x*). If the score is larger than a threshold *τ*, we reject *x* since this implies that we have seen many similar sequences. Otherwise, we store *x* as part of our sample. Algorithm 1 describes our approach. There are several valid string distance measures with corresponding LSH families that are appropriate for this situation, including the Hamming distance, Jaccard distance, and edit distance [19]. In practice, we primarily consider the Jaccard distance over the set of all *k*-mers because the MinHash LSH family [4] can be implemented very efficiently for both genomic and metagenomic applications [23]. However, our results and algorithms generally apply to other string distances.

### 2.2 Computation and Memory Cost

Before we address the intuition and analysis of the RACE diversity sampling algorithm, we review the computation and memory requirements of Algorithm 1. RACE is clearly a one pass streaming algorithm since we only see each element of the dataset once. The computational cost of RACE diversity sampling consists of three basic operations. We need to compute *R* hashes of the sequence string followed by *R* counter look-ups and *R* scalar additions to the count value. In our experiments, we show that we can use *R* as small as 10. Clearly, the computational cost is negligible.

The memory footprint is also small. Unlike reservoir sampling, sketches or coresets, RACE does not need to store buffers of sequence data. Instead, we store a *B × R* array of integers in RAM. Even for very large datasets, we only need *B* = 100*k* and *R* = 10. This is only one million numbers to store, or a few megabytes of RAM.

### 2.3 Intuition

The RACE score is a kernel density estimate 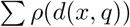 over the sequence dataset, where the distance between sequences is defined as the Jaccard similarity. Each RACE count is a noisy indicator of the number of nearby sequences. Since the RACE estimates sharply concentrate around the kernel density, only a few repetitions are needed to provide a good approximation. If a sequence is in a densely populated region of the dataset, many of the RACE counts will be large. Algorithm 1 downsamples such regions of space. This improves the sequence diversity of the sample since there are many common sequences that do not need to be represented more than once. On the other hand, our algorithm keeps unique sequences because they are likely to have a low score. RACE decides whether to keep a sequence based on the sum of Jaccard similarity scores for each sequence in the sample. We expect RACE to perform better than random sampling because random sampling over-represents densely populated regions and under-represents sparse regions, while RACE attempts to represent both equally.

## 3 Theoretical Results

In this section we present theoretical results which show that Algorithm 1 maximizes the diversity index of the sample. Our proof sketch is as follows. We start with the assumption that the sequence classes are separable using a string distance. Using this assumption, we show that the distribution of the RACE counters for S converges to a uniform distribution under Algorithm 1. We conclude our analysis with a proof that if a sample has uniform RACE counters, then it has an optimal diversity index. We begin by defining the diversity sampling problem.

### Definition 2

*Diversity Sampling Problem*

*Given a streaming dataset* 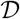 *consisting of elements x*_1_, *x*_2_, …, *x_N_ where each element x_i_ belongs to one class C*_1_, *C*_2_ … *C_S_ and a diversity measure* 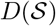, *construct a subset* 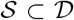 *of M samples, where M* ≪ *N, such that*

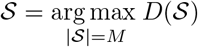

Definition 2 is also given in [11] and the problem has been analyzed for a dozen different diversity measures 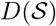. However, all of these measures are low-level geometric functions that do not consider the class labels. The only exception is topic diversity, where we wish to solve a set coverage problem on the topics covered by a sample of news articles. Here, we adopt more traditional biodiversity measures such as the Simpson, Shannon, and Berger–Parker indices. In practice, our classes may be operational taxonomic units (OTU), species, genera or any other classification. We want a subset that represents as many classes as possible.

### 3.1 Convergence of RACE Counts

#### Definition 3.

*Δ-separable classes: We say that two classes C*_1_ *and C*_2_ *are Δ-separable if d*(*x, y*) ≥ *Δ for all x* ∈ *C*_1_ *and y* ∈ *C*_2_

Under any reasonable string representation there is a value of *Δ* > 0 such that the sequence classes are *Δ* separable using the string distance. Under this assumption, we can force each labeled class to map to a unique set of buckets in the RACE array. Lemma 1 is a consequence of standard LSH amplification techniques [9]. To prove the lemma, we use the minimum distance *Δ* to establish bounds on *β* from Definition 1. We then apply the concatenation trick to bound the probability of hash collisions between sequences from different classes.

#### Lemma 1.

*Suppose that two classes C*_1_ *and C*_2_ *are Δ-separable and that there exists a* (*R, Δ,α, β*)-*sensitive hash family* 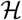. *Then given δ* ∈ [0, 1] *there exists a LSH function h*(*x*) *such that h*(*x*) ≠ *h*(*y*) *for all x* ∈ *C*_1_ *and y* ∈ *C*_2_ *with probability* 1 − *δ*.

*Proof.* Construct a hash function *h*(*x*) by concatenating *n* hash functions sampled from 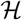 and note that *h*(*x*) is (*R, Δ, α, β*)-sensitive. We want to lower bound the probability that there are no collisions between *x* ∈ *C*_1_ and *y* ∈ *C*_2_.

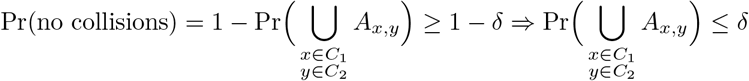

where *A_x, y_* is the event that *h*(*x*) = *h*(*y*). Using the union bound and the fact that *C*_1_ and *C*_2_ are *Δ*-separable, we have that

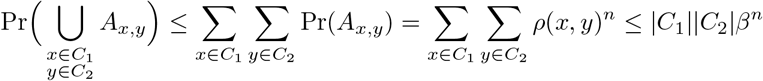

because *d*(*x, y*) ≥ *Δ* for all *x* ∈ *C*_1_ and *y* ∈ *C*_2_. To get the result, put

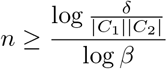

The Kullback-Leibler (KL) divergence is often employed to characterize the relative entropy between two probability distributions. By applying Lemma 1, we can show that the KL divergence between the uniform distribution and the empirical distribution of RACE counts is minimized (i.e., the distribution of the RACE count values converges to a uniform distribution).

#### Theorem 2.

*RACE Count Convergence*

*Construct a single RACE array with B buckets using a LSH function h*(·) *on the sample set* 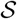 *from Algorithm 1. Then Algorithm 1 causes the distribution of the RACE counters in this array to converge to the uniform distribution.*

*Proof.* Let 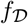 be the underlying probability distribution that generates each element of the data stream 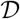. Restrict the domain of *h*(*x*) to 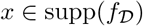 and assume without loss of generality that the restricted range of *h*(*x*) is the set of integers {1…*B*}. Now run Algorithm 1 until all of the *B* counters are nonzero. We will show that the KL divergence between the distribution of RACE counts and the uniform distribution converges to zero. The KL divergence is defined as

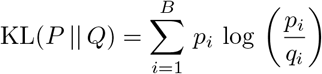

To simplify the analysis, we assume log base 2. Let *Q* be the uniform distribution and let *P_n_* be the distribution of RACE counters after *n* elements have been read from the data stream. Let *Δ_n_* be the divergence between *Q* and *P_n_*. Then

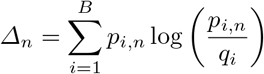

Here *q_i_* = 1/*B* and *p_i,n_* = *b_i,n_*/*m_n_*, where *b_i,n_* is the count value in bucket i and *m_n_* is the sum of all counts at time *n*. The divergence *Δ_n_* is a random variable over the randomness of the hash function and data stream. We want to show that *Δ_n_* converges almost surely to 0, or equivalently that

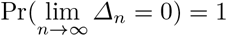

where

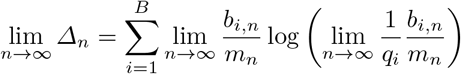

Observe that *b_i,n_* and *m_n_* are non-decreasing bounded sequences. While this alone does not guarantee convergence, it is easy to see from Algorithm 1 that *b_i,n_* attains its supremum ⌊*τ*⌋ and therefore *b_i, n_* → ⌊*τ*⌋ and *m_n_* → *B*⌊*τ*⌋ since 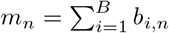. Therefore we have that

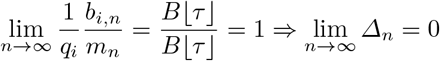

This proves that *Δ_n_* converges almost surely to zero.

### 3.2 Analysis of the Diversity Index

*Diversity Indices* We measure sample diversity using the Shannon index (1), inverse Simpson index (2), and Berger–Parker index (3).

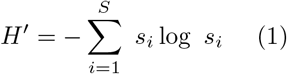

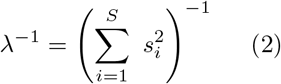

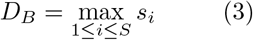

*S* is the number of OTUs, species or, more generally, classes in the sample 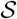. The proportion *s_i_* is the ratio *n_i_*/*m*, where *n_i_* is the number of times that class *i* appears in the sample and *m* is the total sample size. There is a strong connection between Theorem 2 and the KL divergence argument in Section 3.1 and the diversity index. Assume the classes are *Δ*-separable and use Lemma 1 to ensure that each class maps to a unique set of buckets in the RACE array. We can now express the diversity index in terms of the RACE counts.

#### Corollary 1.

*Assume that classes C*_1_, … *C_s_ are Δ-separable and use the hash function from Lemma 1 to construct a RACE array on the output of Algorithm 1. Then the ratio s_i_ converges almost surely:*

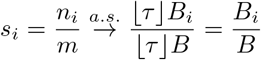

*where B_i_ is the number of buckets in the hash range of class C_i_*.

#### A simple case

Consider the simple case where each class maps to a single bucket (*B_i_* = 1). With *B* = *S* buckets, we have that the inverse Shannon index converges to the maximum value of log *S*. An elegant proof begins with the observation that there is a relationship between *H′* and the KL divergence from Theorem 2.

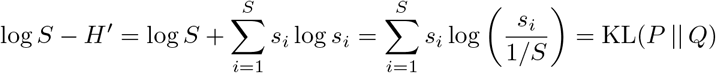

The last equality is obtained using the fact that *s_i_* = *b_i_*/*m* when we assume that *B_i_* = 1 and *B* = *S*. Since KL(*P* || *Q*) → 0, *H′* → log *S*. The key component of this analysis is that *s_i_* → 1/*S*. It is easy to see that this value of *s_i_* also maximizes the other diversity indices. In fact, RACE works for any diversity measure 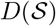 that attains its optimal value when *s_i_* = 1/*S*.

#### The general case

The previous argument assumed that *B_i_* = 1 but does not hold when *B_i_* ≠ 1. However, RACE still converges to the optimal diversity index when each class maps to the same number of buckets. To see this, suppose that each class maps to *k* different buckets. Then Corollary 1 holds with *B_i_* = *k* and *B* = *kS*. Therefore, we have that *s_i_* → *k/kS* = 1/*S*. A number of useful data models have this property. For instance, we might consider a dataset where class *C_i_* consists of a uniform sample on the ball 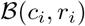. If all the ball radii {*r_i_*, … *r_S_*} are the same, then all the classes will all map to the same number of buckets. This model assumes that the sequences in each class are highly similar to each other, but one can construct other models with the same convergence behavior that do not require this assumption.

Although useful in theory, all of our assumptions are too restrictive in practice. Classes often do not map to the same number of buckets. However, Corollary 1 still provides strong convergence guarantees and useful intuition. If each class maps to *approximately* the same number of buckets, then RACE sampling will produce a sample with a diversity index that is nearly optimal. Furthermore, Corollary 1 indicates that we will oversample classes with high sequence diversity. OTUs with large pan-genome sizes will cover a larger number of buckets in the RACE array than OTUs with small pan-genomes. For instance, RACE sampling would likely retain many samples belonging to *Escherichia coli* bacteria in metagenomics applications. *Escherichia coli* has an open pan-genome with an incredibly high degree of sequence diversity [10]. Therefore, *Escherichia coli* will cover a larger number of RACE buckets and have a higher *B_i_* than a species with a smaller, closed pan-genome. In general, Corollary 1 shows that RACE sampling attempts to preserve sequence diversity based on the Jaccard string metric. RACE will oversample classes that cover large volumes of the Jaccard metric space. Equivalently, RACE oversamples OTUs with large pan-genomes but attempts to sample from each OTU present in the sample. In contrast, random sampling will oversample common OTUs with large populations. Therefore, our theory shows that RACE is an effective method for rare-species sampling in metagenomic studies.

## 4 Experimental Results

The goal of this section is to evaluate the diversities of the sample sets produced by RACE against baseline sampling methods. We use a variety of methods to produce the same number of samples and evaluate them based on the number of taxa represented in the subsample and the Shannon Diversity Index. We also aim to provide a characterization of the time needed to run our method against competing baselines.

### 4.1 Datasets

We use metagenomic microbiological studies accessed through the ENA to evaluate our methods. We found that metagenomic datasets tend to have separable classes with reasonable sequence diversity. Consequently, sequences belonging to the same species often have much higher Jaccard similarities compared to sequences belonging to different species. It should be noted that FASTQ files from the ENA do not specify the species that corresponds to each each sequence. To obtain the ground truth reference labels we used Kraken2 to classify each read with a taxon label. While relying on Kraken2 to label metagenomic reads has been previously shown to be both sensitive and accurate, this process can induce bias if applied to microbiomes with poor representation in the Kraken2 database [22]. To address this, we focused on human-host associated microbiomes from the HMP2 project [24], where we found that over 70-80% of the microbial species in these samples were contained in the database. Before performing the classification and sub-sampling, we used Trimmomatic [3] to remove errors in the read data. Additional information is available in Table 1.

**Table 1.** Dataset information. We report the ENA run accession, dataset size *N*, mean sequence length *L,* number of species in the dataset *S*, and a description of the application. All of our datasets are for Illumina HiSeq 2000 data.

| Dataset | $N$ | $L$ | $S$ | Description |
| --- | --- | --- | --- | --- |
| SRR4343843 | 81,870 | 456.8 | 849 | Odor-causing bacteria in automobile air conditioning systems |
| SRR1565331 | 269,247 | 71.0 | 1120 | HMP2 sample of anterior nares |
| SRR1804823 | 9,199,019 | 98.6 | 6387 | HMP2 sample of supragingival plaque |
| SRR2175724 | 10,214,210 | 98.4 | 6759 | HMP2 sample of fecal matter |
| SRR1804468 | 501,590 | 99.4 | 3630 | HMP2 sample of buccal mucosa |

### 4.2 Experiment Setup

We implemented random sampling, composable coresets and used diginorm as a baseline. We briefly describe each of our methods:

1. **Composeable coresets:**We used the Jaccard distance and a window length of 100 sequences.
2. **Reservoir Sampling:** We used the default reservoir sampling method that stores a buffer of sampled elements.
3. **Diginorm:** We used the khmer software package with the default Diginorm settings [8]. To show results for different sample sizes, we varied the Diginorm threshold.
4. **RACE:** We varied *τ* to represent a wide range of subsample sizes.

*RACE Array Parameters* The RACE algorithm requires five parameters: *τ*, *R*, *B*, *k*, and *n*. The size of the RACE arrays is determined by *R* and *B*, while the number of samples in 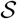 is related to *τ*. *R* is the number of RACE repetitions. *B* is the number of hash buckets in each array of counters. We use the rehashing trick to bound the range of the hash function to the interval [1, *B*]. In practice, sequences do not occupy the whole range of each hash function. Thus, we can reduce the memory needed by our method without losing much information by rehashing to a narrower range than predicted by the theory. Although the memory usage of the RACE algorithm is minimal even without rehashing, smaller RACE count arrays can be processed much more quickly because they fit into the CPU cache. Increasing *τ* increases the number of samples that will be saved by RACE. There is no closed-form expression for the number of samples that RACE will accept for a given *τ*. However, a good heuristic is to set *τ* to be approximately equal to the desired number of sequences from each species in 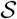. In the experiments, we used *R* = 10 and *B* = 100, 000. However, the algorithm does not seem to be sensitive to these parameters. Also note that for very small values of *τ*, we increased *R* to 50 or 100 to be able to produce smaller sketches.

*LSH Parameters* We use MinHash on all *k*-mers in the short read sequence. To construct the LSH functions, we need to choose *k* and *n*, the number of LSH concatenations. *k* is the number of base pairs in the *k*-mers used to vectorize the sequence. The *k* parameter controls the Jaccard similarity score between sequences and *n* controls the sensitivity of the algorithm. Increasing *k* makes it less likely for two sequences to collide under MinHash. The decrease in collision probability is larger for sequences from different OTUs than for sequences in the same OTU. We characterized this behavior for one of our datasets (Figure 2) to show how *Δ*-separablility depends on *k*. In general, larger values of *k* and *n* are required to separate highly similar sequences. RACE makes sampling decisions based on the Jaccard similarity scores determined by *k* and *n*. To convert similarity score thresholds to standard alignment-based measures such as the average nucleotide identity (ANI), one can use the MASH distance [23] or the more in-depth ANI estimation procedure proposed by FastANI [13]. When *n* = 1, the guidelines for choosing *k* from [23] and [13] directly apply to the RACE algorithm. For the human microbiome datasets, we used *k* = 18 and *n* = 1. For the additional experiments in the Appendix, we used *k* = 6 and *n* = 4.

**Fig. 1.**
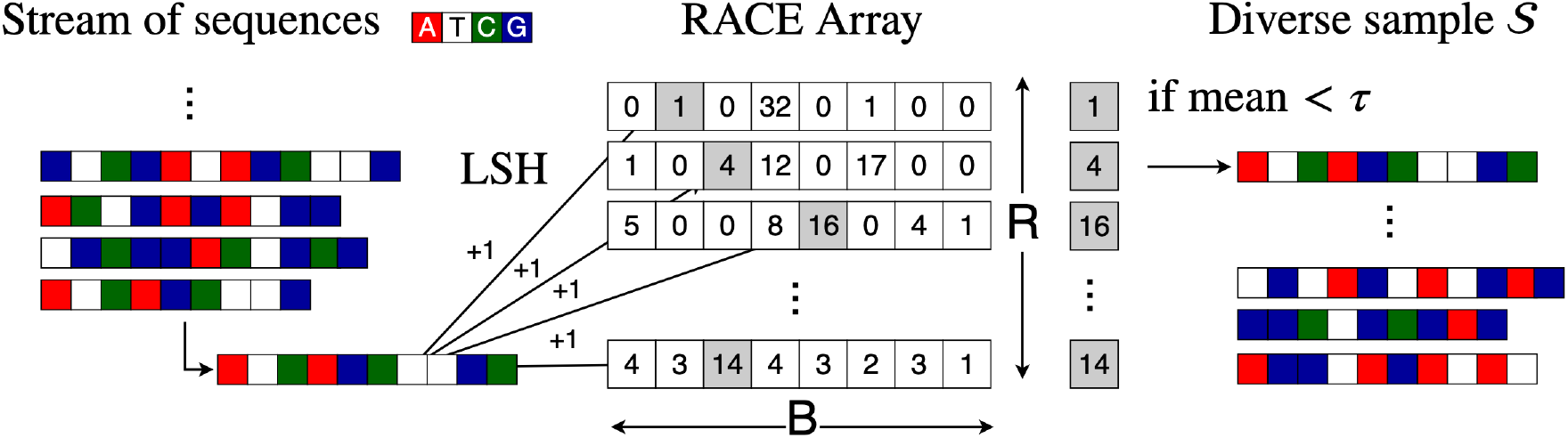
Illustration of the RACE diversity sampling algorithm (Algorithm 1). To create a diverse sample 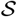 of a sequence data stream, we hash each element with *R* LSH functions and find the mean (score) of the corresponding array locations and increment them. If the mean is smaller than *τ*, we keep the sequence. Otherwise, we discard it.

**Fig. 2.**
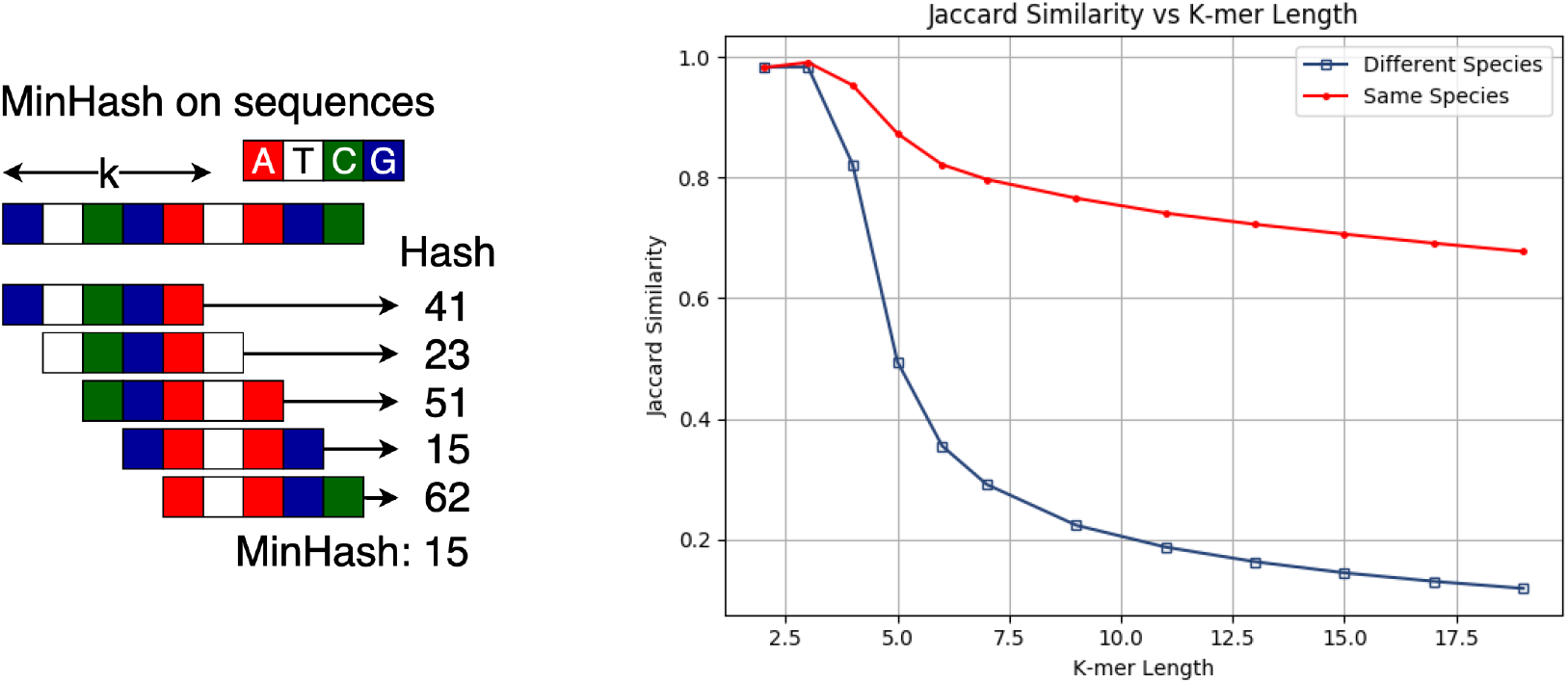
MinHash on short read DNA sequences. Based on the *k*-mer length, different taxa (clusters) of sequences may be easy or difficult to separate using MinHash. The collision probability of MinHash (left) is equal to the Jaccard similarity (right).

### 4.3 Results

We provide two sets of experiments. On our smaller datasets, we provide an exhaustive comparison with Diginorm and coreset baselines in Figure 3. However, coresets have a very low throughput and Diginorm requires > 15GB of RAM for large datasets. Therefore, we only show comparisons between RACE and random sampling at this scale (Figure 4). We also show that RACE preserves the Shannon diversity index in addition to sampling more species (Figure 5). We show how sequence similarity varies with k-mer length in Figure 2.

**Fig. 3.**
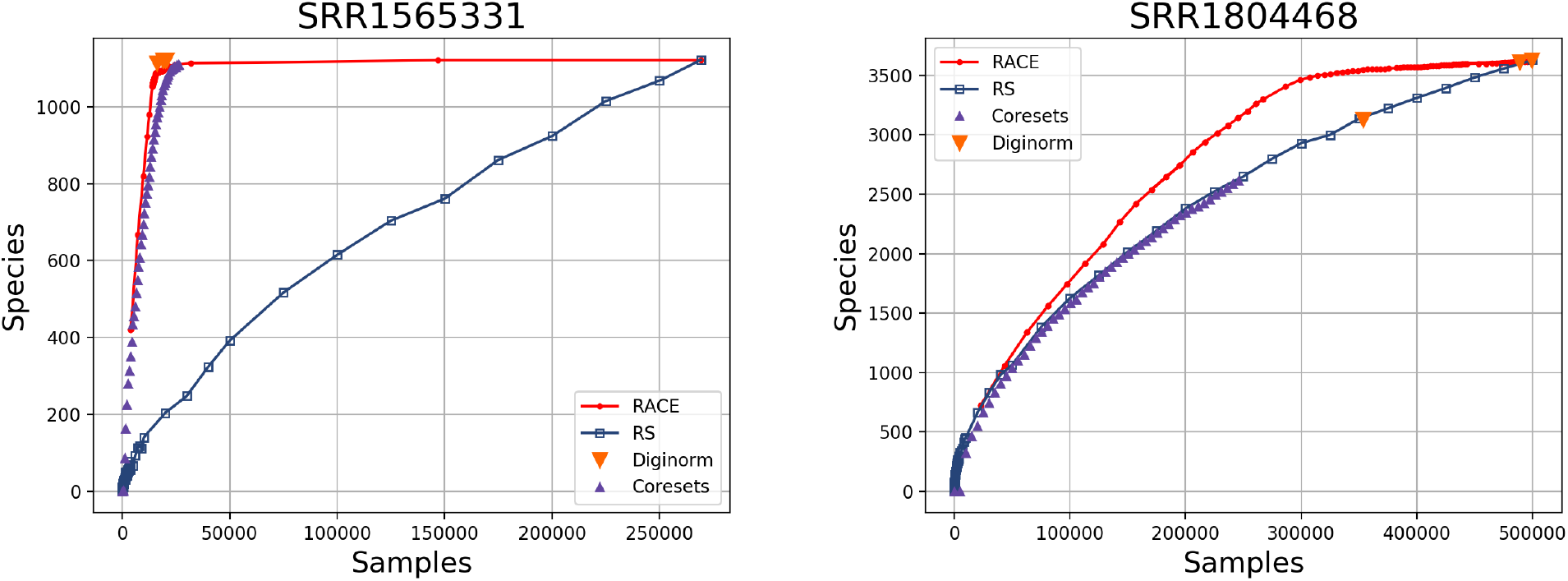
Number of species represented in a sample for RACE, random sampling (RS), composeable coresets and Diginorm. Based on the Diginorm thresholding process, we were unable to specify the full range of sample sizes.

**Fig. 4.**
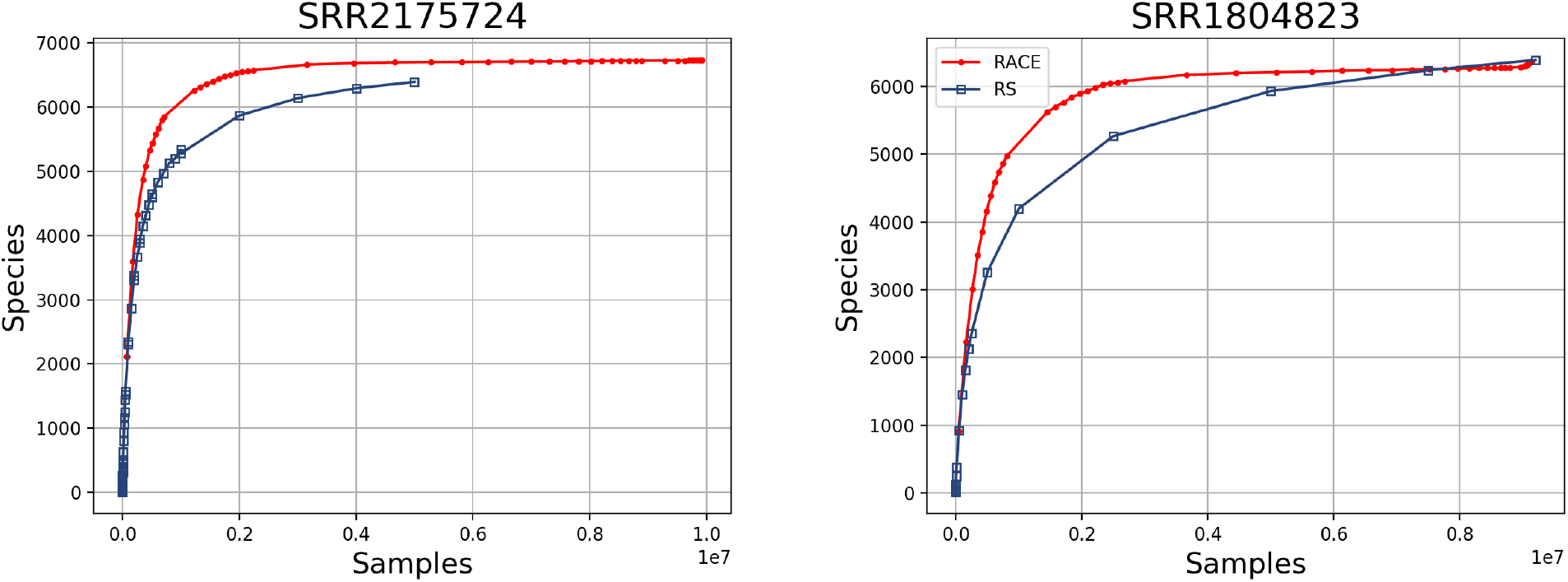
Experimental results for large-scale datasets (> 10^7^ reads). Due to the large scale of the datasets, coresets were computationally infeasible and Diginorm required large volumes of RAM.

**Fig. 5.**
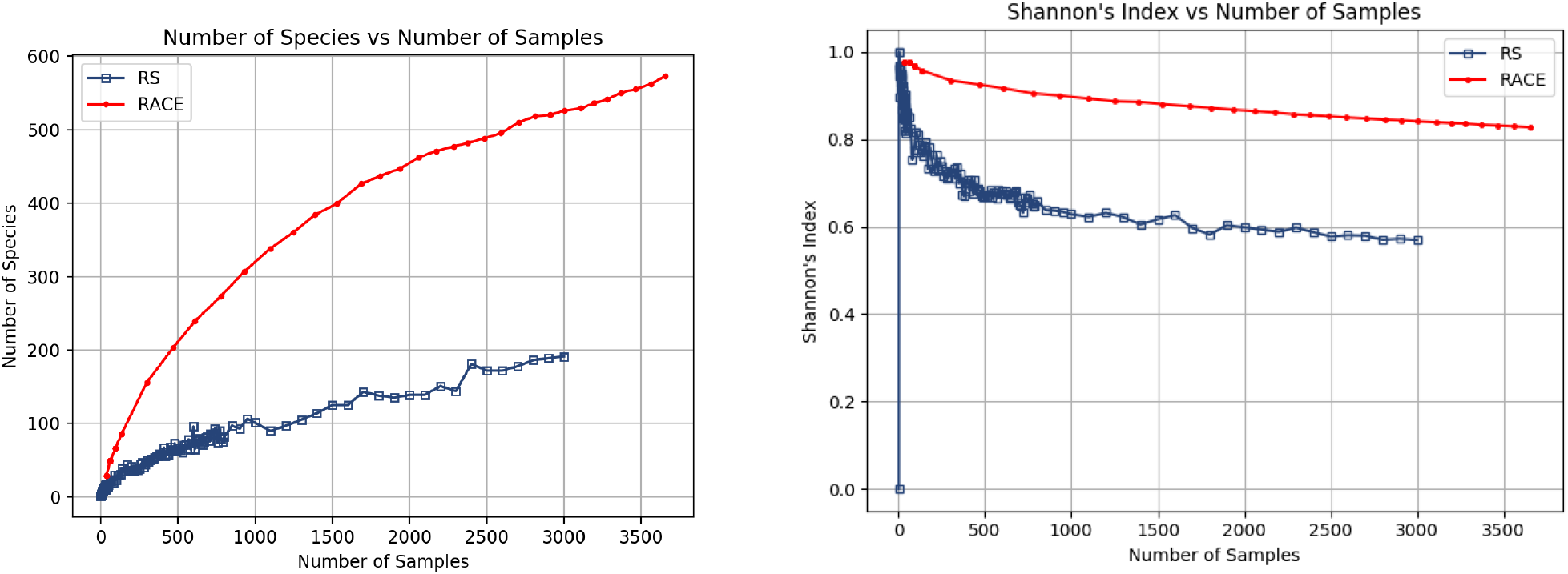
Experimental results on SRR4343843 with τ from 0.1 to 30. We observe that RACE increases both the number of species represented in the sample and the Shannon Index, showing that we do obtain a more diverse sample than our random sampling baseline.

## 5 Discussion

From the results reported above, RACE consistently delivers highly competitive metagenomic diversity compared to the provided baselines. RACE and random sampling were the only two algorithms capable of scaling to large dataset sizes under reasonable computational constraints. While Diginorm and coreset methods can sometimes provide samples with similar diversity to those obtained using RACE, it comes at a high resource cost (Table 2). Coresets are computationally intensive and Diginorm is memory intensive when compared to RACE. We found that Diginorm can consume memory that is upwards of 1000x the amount consumed by RACE and that coresets have 200x lower throughput. While Diginorm and coresets are appropriate for offline summarization and small datasets, neither algorithm provides the reliable and efficient diverse sampling performance we observed using RACE. The large memory difference between RACE and random sampling is due to the fact that random sampling needs to store a reservoir of sequences in RAM. This is required because we need to be able to discard samples from the reservoir. RACE, on the other hand, can immediately output the sequence as it arrives or dump it to disk once accepted. Furthermore, the entire RACE data structure fits in the cache of the CPU, leading to both memory and computational advantages.

**Table 2.** Computational resources comparison. We report the data throughput, RAM usage, and (where available) the 99^th^ percentile latency. The 99^th^ percentile latency is the maximum time needed to process 99% of streaming inputs. It determines the data rate, as only 1% of inputs take longer to process. Note that the memory for random sampling depends on the sample size and the memory for Diginorm depends on the input size.

| Method | Throughput (Mbp/sec) | RAM | Latency |
| --- | --- | --- | --- |
| RACE | 2.6 | 16 kB - 16 MB | 366 μs |
| Diginorm | 6.4 | 500 MB - 15 GB | – |
| Random Sampling | 122 | 50 MB - 2 GB | 7 μs |
| Coresets | 0.012 | 50 MB - 2 GB | 1.8 s |

We expect that RACE will be broadly useful for OTU diversity sampling, species sampling and sampling at higher taxonomic levels. While our thresholds are defined in terms of Jaccard similarity scores, these scores are closely related to more conventional ways to differentiate between genomes such as ANI [23]. By adjusting the similarity thresholds and changing the *k* and *n* parameters of the algorithm, we can make RACE sufficiently sensitive to differentiate between highly similar sequences within the same OTU or sufficiently indiscriminate to make coarser classifications. This utility will make RACE broadly useful. For instance, when combined with read-until hardware technology, RACE may be able to construct the most informative sample from the stream of genetic information.

## 6 Conclusion

We have presented RACE, a streaming algorithm that constructs a diverse sample using minimal memory and computation. Advances in DNA sequencing techniques enable the production of genetic data at a rate that surpasses that at which we store and analyze the data. Our RACE streaming algorithm is a simple and highly effective solution for compressing these massive datasets while preserving metagenomic diversity and species representation. Based on similar speedups from coresets in computational geometry, we expect that RACE will enable analyses with a smaller memory and computational footprint. The simplicity, speed and low memory consumption of RACE compared to similarly-performing baselines provides ample evidence that RACE is an effective method to compress genetic datasets for transmission, processing and storage. Our code is available at https://github.com/brc7/DiversitySampling.

## Supporting information

Supplemental Experiments

## Footnotes

1 Code is available at https://github.com/brc7/DiversitySampling

