## Supplemental Experiments for "Diversified RACE Sampling on Data Streams Applied to Metagenomic Sequence Analysis"

### Diversified Sampling on Data Streams Applied to Sequence Analysis: Supplementary Material

| Label | Dataset | $N$ | $L$ | $S$ | Description |
|---|---|---|---|---|---|
| A | SRR1056036 | 19,431 | 465.9 | 229 | Bacteria in Seawater |
| B | SRR4454042 | 765,139 | 132.4 | 1740 | NGS sequence of V3 region of soil metagenome |
| C | SRR3744867 | 1,636,283 | 344.6 | 3279 | Bacteria community in seawater at 14N |

**Table 1.** Supplementary dataset information. We report the ENA run accession, dataset size $N$, mean sequence length $L$, number of taxa in the dataset $S$ and a description of the application.

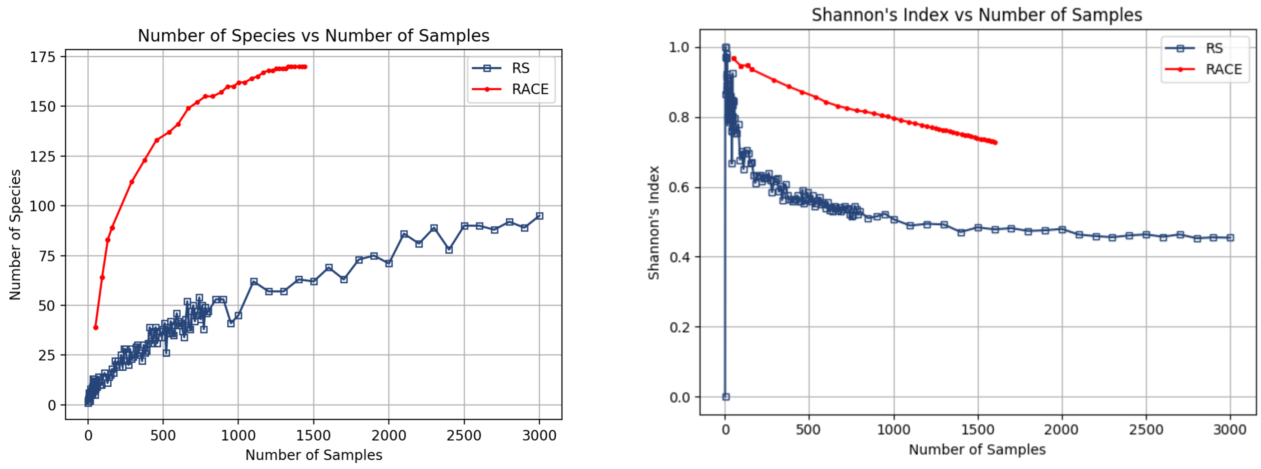


**Fig. 1.** Experimental results on dataset A with $\tau$ from 0.1 to 30

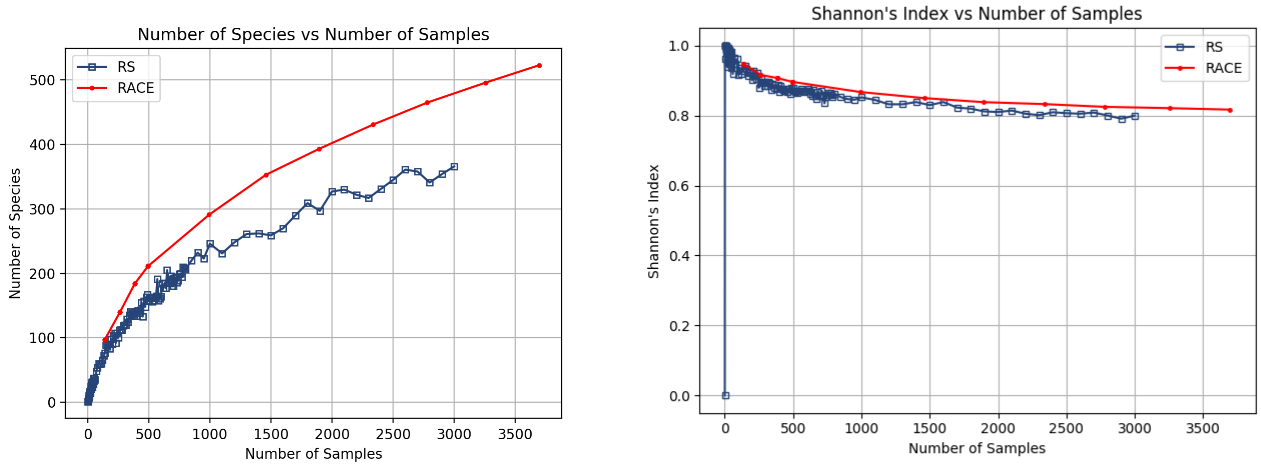


**Fig. 2.** Experimental results on dataset B with $\tau$ from 0.1 to 8

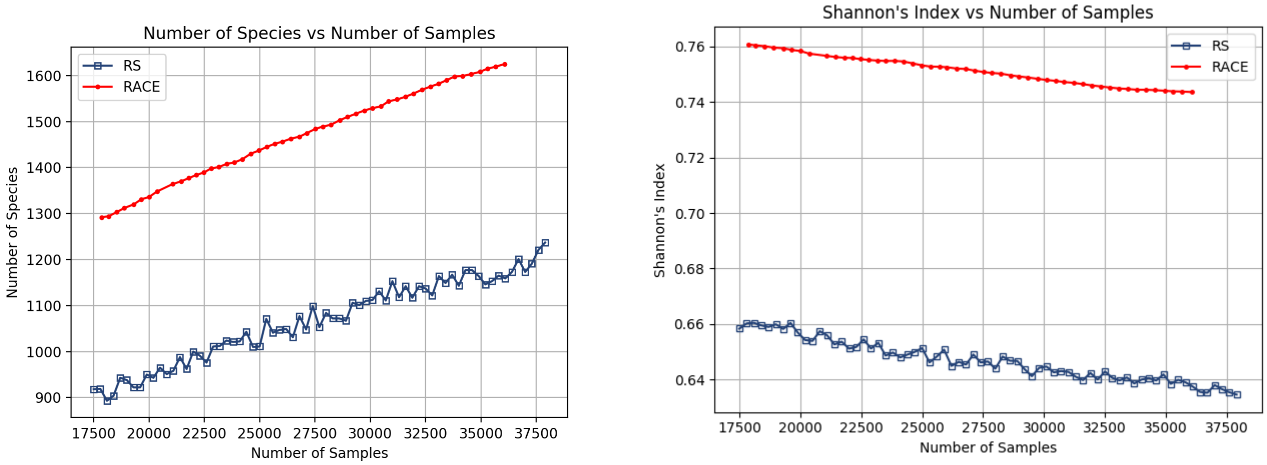


**Fig. 3.** Experimental results on dataset C with $\tau$ from 50 to 100
